# Count does not recover major events of gene flux in real biological data

**DOI:** 10.1101/246272

**Authors:** Nils Kapust, Shijulal Nelson-Sathi, Barbara Schönfeld, Einat Hazkani-Covo, David Bryant, Peter J. Lockhart, Mayo Röttger, Joana C. Xavier, William F. Martin

## Abstract

In prokaryotes, known mechanisms of lateral gene transfer (transformation, transduction, conjugation and gene transfer agents) generate new combinations of genes among chromosomes during evolution. In eukaryotes, whose host lineage is descended from archaea, lateral gene transfer from organelles to the nucleus occurs at endosymbiotic events. Recent genome analyses studying gene distributions have uncovered evidence for sporadic, discontinuous events of gene transfer from bacteria to archaea during evolution. Other studies have used traditional birth-and-death phylogenetic models to investigate prokaryote genome evolution to claim that gene transfer to archaea was continuous during evolution, rather than involving occasional periodic mass gene influx events. Here we test the ability of Count, a birth-and-death based program, to recover known events of mass acquisition and differential loss using plastid genomes and eukaryotic protein families that were acquired from plastids. Count showed a strong bias towards reconstructed histories having gene acquisitions distributed uniformly across the tree. Sometimes as many as nine different acquisitions by plastid DNA were inferred for the same protein family. That is, Count recovered gradual and continuous lateral gene transfer among lineages, even when massive gains followed by gradual differential loss is the true evolutionary process that generated the gene distribution data.

## Introduction

Lateral gene transfer (LGT) has had a major impact on gene distributions among archaeal chromosomes during evolution (Wagner et al. 2017). There are basically two ways to infer the evolutionary processes underlying gene distributions. One approach is to construct phylogenetic trees for all proteins in a given set of genomes and to compare topologies in search of phylogenetic congruence or incongruence, evoking vertical inheritance to account for the former and LGT to account for the latter. Despite the occurrence of historical events of lateral gene transfer among prokaryotes, applications of this approach have nevertheless generally led to phylogenetic reconstructions favoring a single dominant underlying prokaryotic tree (e.g. Daubin et al. 2003). One limitation of this investigative approach, and thus the conclusions evidenced, is that it is hampered by the circumstance that the vast majority of genes in prokaryotes occur in only a very few genomes (Dagan and Martin 2007). Genes present in only two or three genomes will appear to have been vertically inherited in all trees, and ≥ 1/3 of all genes present in four genomes will also appear to be vertically inherited by phylogenetic congruence criteria alone. The problem with this potential methodological bias is that it will inflate ancestral genome sizes to unacceptably large values if one looks at all genes (Dagan and Martin 2007), not just the ones for which trees are convenient to construct.

A different and still relatively new approach to investigate the factors underlying gene distributions is to cluster all protein coding genes in a given set of genomes into protein families and to examine not only the presence and absence patterns (PAPs) of those genes along a given reference tree, but also the phylogenies for each individual cluster (Nelson Sathi et al. 2012; Ku et al. 2015). When applied to archaea, this approach uncovered that haloarchaea acquired about 1000 genes from bacteria in a process that transformed a chemolithoautotrophic methanogen ancestor into a facultative aerobic heterotroph (Nelson-Sathi et al. 2012) and that gene acquisitions from bacteria followed by extensive differential loss was important in the origin and evolution of several major archaeal clades (Nelson-Sathi et al. 2015). The same fundamental pattern is observed in eukaryote evolution, where the host lineage is thought to descend from archaea (Martin and Müller 1998; Williams et al. 2013; McInerney et al. 2014; Zaremba-Niedzwiedzka et al. 2017), namely events of mass gene acquisition followed by differential loss (Ku et al. 2015), which is increasing considered a very important factor in genome evolution (Albalat and Cañestro 2016).

Yet another approach to understand gene distributions is to try to reconcile all topologies, all gene duplications, all gene losses, and all gene transfers simultaneously from a given data set (Szöllősi et al. 2015a). The trouble with this approach is that the number of parameters in such a model becomes very large, and there is the risk of overparameterization of models and of falling prey to statistical artefacts, as was recently observed for analyses of gene phylogenies addressing mitochondrial origin (Martin et al. 2017a).

Recently, Groussin et al. (2016) reanalyzed the data of Nelson-Sathi et al. (2015) using a program called Count (Csűrös, 2010). Count takes a given set of PAPs that is determined independently of a reference tree and distributes them across the reference tree allowing LGT and losses (birth-and-death) according to pre-specified parameters that correspond to settings in the Count software. Groussin et al. found basically the same amount of LGT as Nelson-Sathi et al. (2015) found, but Count distributed the LGTs across the reference tree in such a way as to evenly distribute gains and losses according to the settings of the Count program. From that result, they concluded that LGT was mostly uniform and continuous during archaeal evolution (Groussin et al. 2016), not episodic (Nelson-Sathi et al. 2015). However, the same Count method also infers vast amounts of continuous LGT during eukaryote evolution (Szöllősi et al. 2015b), even though there are no known genetic mechanisms for LGT among eukaryotes (Martin 2017), in contrast to the very well characterized mechanisms of LGT among prokaryotes (Popa and Dagan 2011). There are reasons to suspect that the amounts of LGT that Szöllősi et al. (2015b) found for fungi (eukaryotes) are methodological artefacts, because if eukaryotes were exchanging genes freely across higher taxonomical boundaries then eukaryote genomes should exhibit cumulative effects of LGT as prokaryote genomes do, but the converse is observed (Martin 2017). Moreover, genome-scale tests for eukaryote LGT show that gene evolution in eukaryotes is vertical, mediated by loss and punctuated by gene acquisitions at endosymbiotic events (Ku et al. 2015; Ku and Martin 2016).

Count makes a large number of simplifying assumptions, and we suspect that these modelling assumptions could be responsible for the unusual results returned by the software. The most critical assumption in this context is that the evolutionary histories of different gene families are independent of one another. Thus, an LGT involving a transfer of x genes would be considered as x individual events. Major acquisition events fall completely outside the scope of the model. To examine the impact of this model misspecification, and to test whether it can indeed mislead analyses, we inspect the results produced by Count on real data that evolved by a loss only model, namely chloroplast genomes, to see whether it infers LGT instead of the true process (loss only). We also investigate two other datasets involving gene acquisitions via endosymbiosis to see how Count performs.

## Materials and Methods

### Data collection and annotation

#### Archaeal protein families

The dataset used for the study of the origin of archaeal protein families included 1,981 prokaryotic genomes - 134 archaea and 1,847 bacteria (Nelson-Sathi et al. 2015), hereafter referred to as AR dataset. The amino acid sequences were retrieved from RefSeq, NCBI (version June 2012). The dataset consists of 254,938 archaeal proteins in 25,762 protein families, of which the subset consisting of the import clusters (13,631 archaeal proteins in 2,264 protein families), used in Groussin et al. (2016), was used as well here.

#### Plastid protein families

A dataset encompassing all plastid encoded proteins for 193 photosynthetic eukaryotes (Schönfeld 2012), designated as the PL dataset, was used. It consists of 254 protein families from 193 sequenced plastid genomes of different eukaryotes, encompassing 6561 protein sequences in total. All sequences were retrieved from RefSeq, NCBI (version January 2011). Each protein family was manually annotated into Uniprot functional categories.

#### Eukaryote protein families

The eukaryotic protein dataset was taken from Ku et al. (2015), hereafter referred to as the EK dataset. It contains 21,146 protein sequences from 55 eukaryotic genomes from six different supergroups. The dataset was divided into two different matrices: one for 1,060 protein families shared in photosynthetic eukaryotes and densely distributed in cyanobacteria (6528 sequences, corresponding to block A, B and C in Ku et al. (2015)) and another for 1,397 protein families present in the eukaryotic common ancestor that are likely to correspond to the origin of the mitochondrion (14,618 sequences corresponding to block E in Ku et al. (2015)).

For each dataset, a PAP was constructed. In the PAPs, each row corresponds to a species and each column to a protein family, binary elements of the matrix indicate presence or absence in the respective genome. Phylogenetic reference trees for the AR and EK datasets were taken from Nelson-Sathi et al. (2015) and Ku et al. (2015) respectively. For the PL dataset, the reference tree was assembled from Schönfeld (2012) based on Bayesian inference of trees for the individual genes. Internal nodes are designated as HTUs (hypothetical taxonomic units), terminal nodes as OTUs (operational taxonomic units).

### BLAST against cyanobacterial genomes

The 15,588 protein sequences in the PL dataset were blasted against 94 cyanobacterial genomes retrieved from RefSeq, NCBI (version September 2016, listed in Supplemental Table 1). Hits were filtered with a threshold of e-value equal to or less than 1e-10 and local identity equal to or greater than 25%.

### Calculation of gain and loss events with Count

Version 10.04 of Count (Csűrös, 2010), written in Java, was used. As input, Count requires a PAP and the corresponding phylogenetic reference tree. Count’s three methods for the analysis of gene evolution – two methods of maximum parsimony, Dollo (DP) and Wagner (WP) and the phylogenetic birth-and-death model (BD) – were tested. The reference tree and the appropriate PAP were loaded into Count (branch lengths are ignored in parsimony models and were not used for the BD model). The data was then optimized using likelihood, a necessary step in order to use the birth-and-death model. All model parameters used were the default Count parameters (Groussin et al. 2016). The following settings were used: the model type was the gain-loss type, the family size distribution at the root was set to Poisson, lineage-specific variation was left unspecified, the gain variation across families was set to 1 for the edge length, the loss and the gain rate. The maximum number of optimization rounds was set to 100 with a convergence threshold on the likelihood of 0.1. The results of the different methods were displayed for each Count record in the graphical user interface, and then evaluated using a Perl script. The respective phylogenetic trees were processed and the results were recorded.

Trees were drawn with FigTree from the results provided by Count. The gain and loss events of the protein families for the respective method were summed and mapped for each corresponding node, respectively, in the phylogenetic tree. For the phylogenetic birth-and-death model the computed numbers for each protein family were rounded up (≥ 0.5) and down (< 0.5) respectively.

## Results

### Reproducing Count’s results for the origin of archaeal protein families

To reproduce the result of Groussin et al. (2016), we analyzed the subset of the AR dataset (Nelson-Sathi et al. 2015) that they analyzed using the phylogenetic birth-and-death model of Count. A comparison (Supplemental Figure 1) shows that the number of gains calculated here using Count vs gains calculated using Count in Groussin et al. (2016) differed only very slightly and only for two archaeal groups (Thermococcales - 58 vs 56 - and Haloarchaea - 219 vs 215). The reasons why Count produced very slight differences for six out of 568 gain events at the roots of the groups in our hands vs. the results of Groussin et al. (2016) are not quite clear but they are also neither cause for concern nor the focus of our interest.

More important is the circumstance that Count attributed no gains to the root of the archaeal tree in our analyses, nor did it do so in Groussin et al. (2016). Supplemental Figure 1b shows the number of different origins per archaeal protein family calculated here for the AR dataset. For 1,726 of the 2,264 archaeal protein families analyzed, Count calculated a single gain event, for 451 protein families two different origin events, for 87 families three different origins and for four of the protein families 6 different origins. For none of the protein families did Count calculate an origin at the root of the archaeal reference tree (Groussin et al. 2016).

### Count does not recover a loss only process

To see whether Count can recover even an obvious process of massive gain followed by differential loss, we examined plastid genomes. It is generally accepted that plastids arose from cyanobacteria via endosymbiosis (Schwartz and Dayhoff 1978). It is also generally accepted that plastid genomes underwent reduction during evolution (Ohyama et al. 1986), that many genes were transferred to the nucleus during evolution and that many gene losses from cpDNA occurred in independent lineages (Martin et al. 1998; Martin et al. 2002). Figure 1 shows the PAPs for chloroplast encoded proteins in a sample of photosynthetic eukaryotes. A BLAST search against 94 cyanobacterial genomes (Supplemental Table 1) shows that 95% of the sequences (highlighted in Supplemental Figure 2) have readily identifiable homologs in cyanobacteria. The tree is rooted with *Cyanophora*, but other roots, including the red lineage have been proposed (Rodríguez-Ezpeleta et al. 2005).

**Fig. 1:**
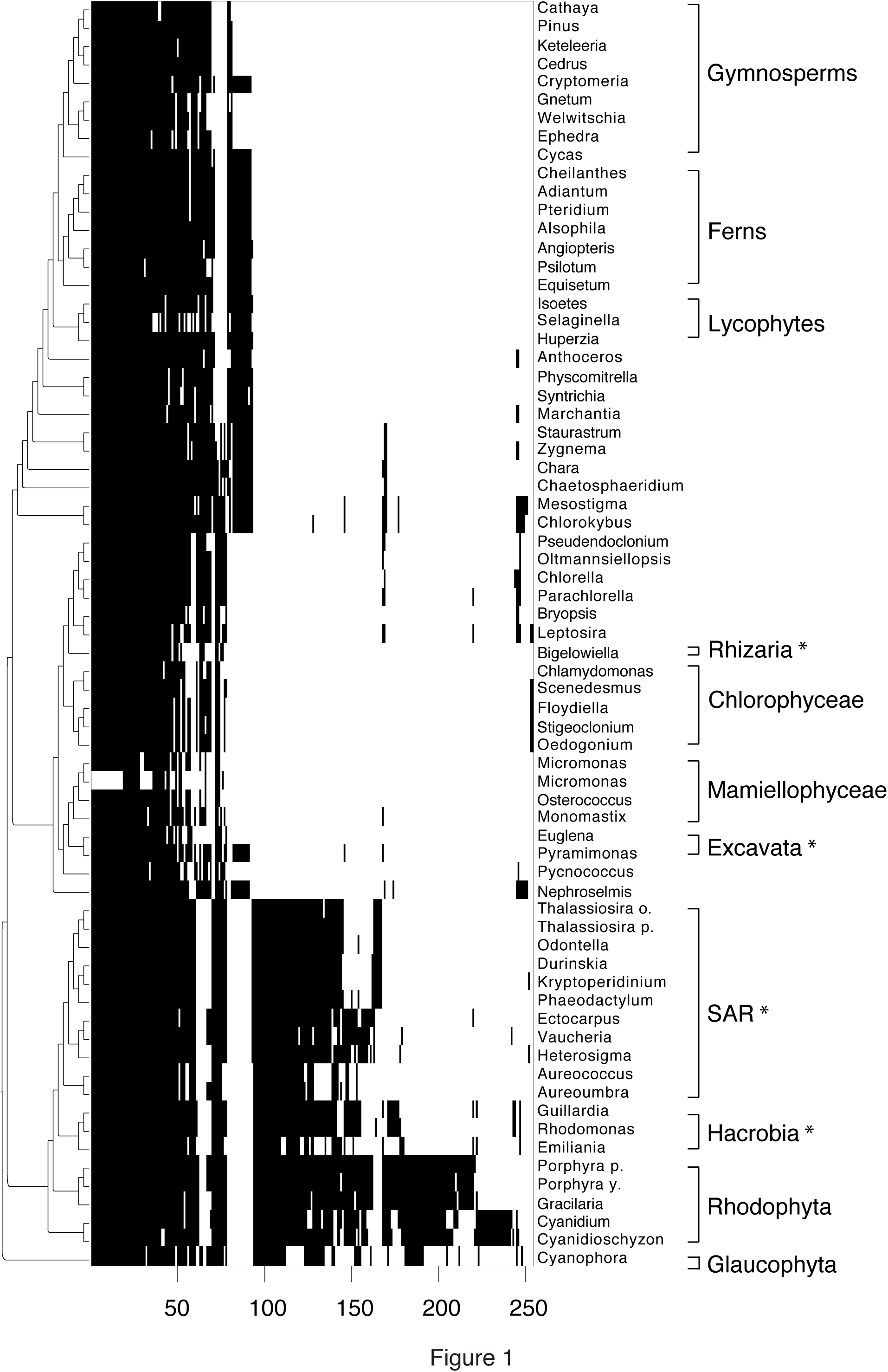
Presence-absence pattern of plastid protein families of the PL dataset. Each black tick indicates the presence of a protein in an OTU. The number of protein families is indicated on the x axis. On the right side of the matrix are the OTUs, on the left the corresponding phylogenetic reference tree. Groups containing secondary plastids are marked with an *.

Regardless of whether we use the parsimony or the birth-and-death options of Count, the program only counts about half of the 254 protein families as being present in the plastid ancestor (Figure 2 and Figure 3a). The other half of the (n.b.) plastid-encoded proteins are reconstructed by Count to have been acquired after the initial plastid, during plant evolution. That is, Count indicated that the primary endosymbiotic event involved acquisition of half a plastid followed by later aquisition of the other half via LGT events in independent lineages. In the birth-and-death model that Groussin et al. (2016) used, Count reports that 86 protein coding genes were acquired once and 36 protein coding genes were acquired twice in the process of lineage diversification during plastid evolution. That is, Count calculates that those 122 genes were acquired from cyanobacteria after lineage divergence during plant evolution and then laterally transferred among eukaryotes. Count does not specify donor or recipient lineages. Another five protein coding genes were acquired three times during plastid evolution.

**Fig. 2:**
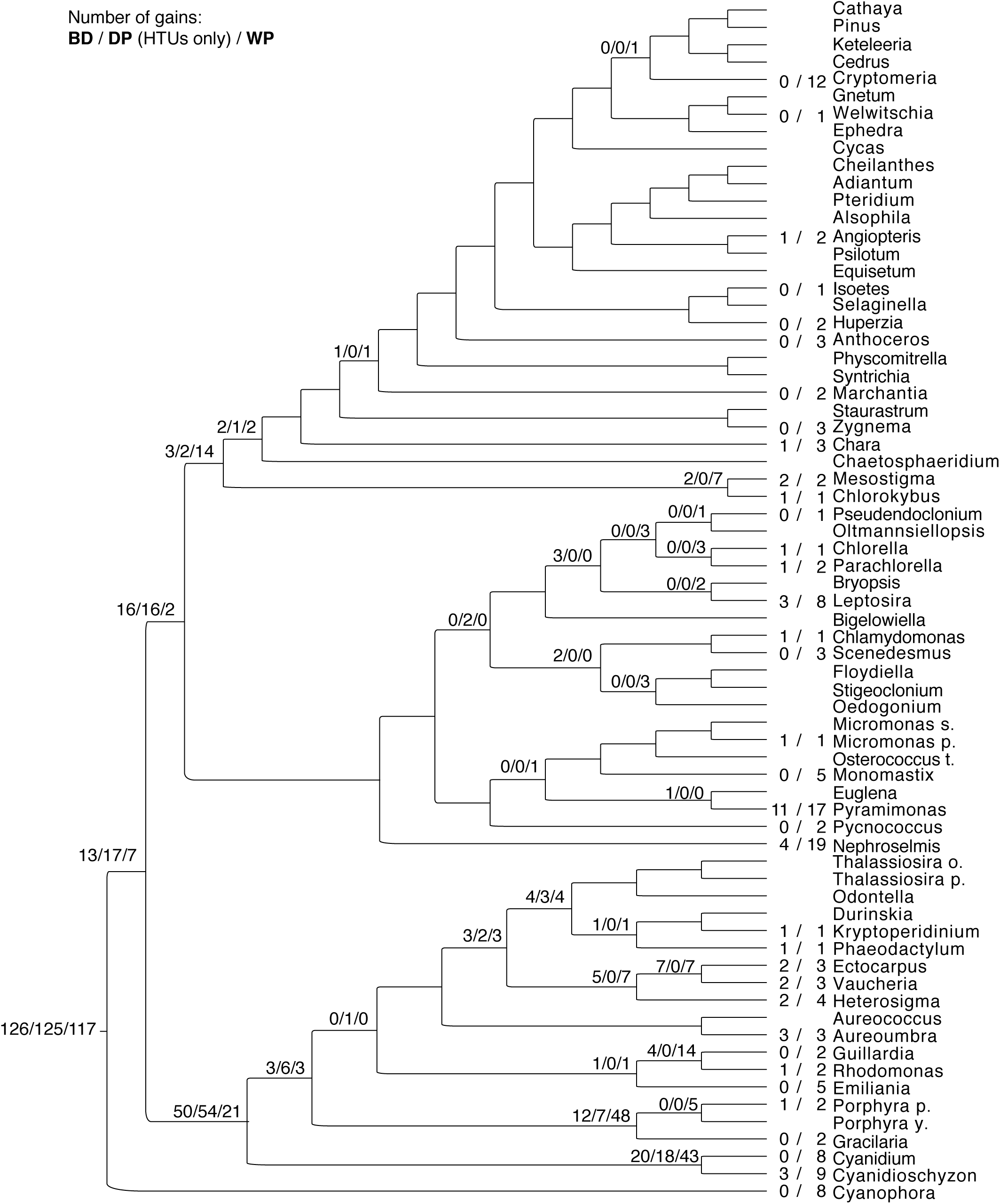
Phylogenetic reference tree for the PL dataset with mapped gain events calculated with Count’s traditional phylogenetic methods. Gain events for plastid protein families are depicted at the respective nodes in the following order, separated by slashes: Birth-and-Death model; Dollo Parsimony (only in the inner nodes); Wagner Parsimony. Inner and outer nodes where no values are plotted have no gain events according to the calculations of Count.

**Fig. 3:**
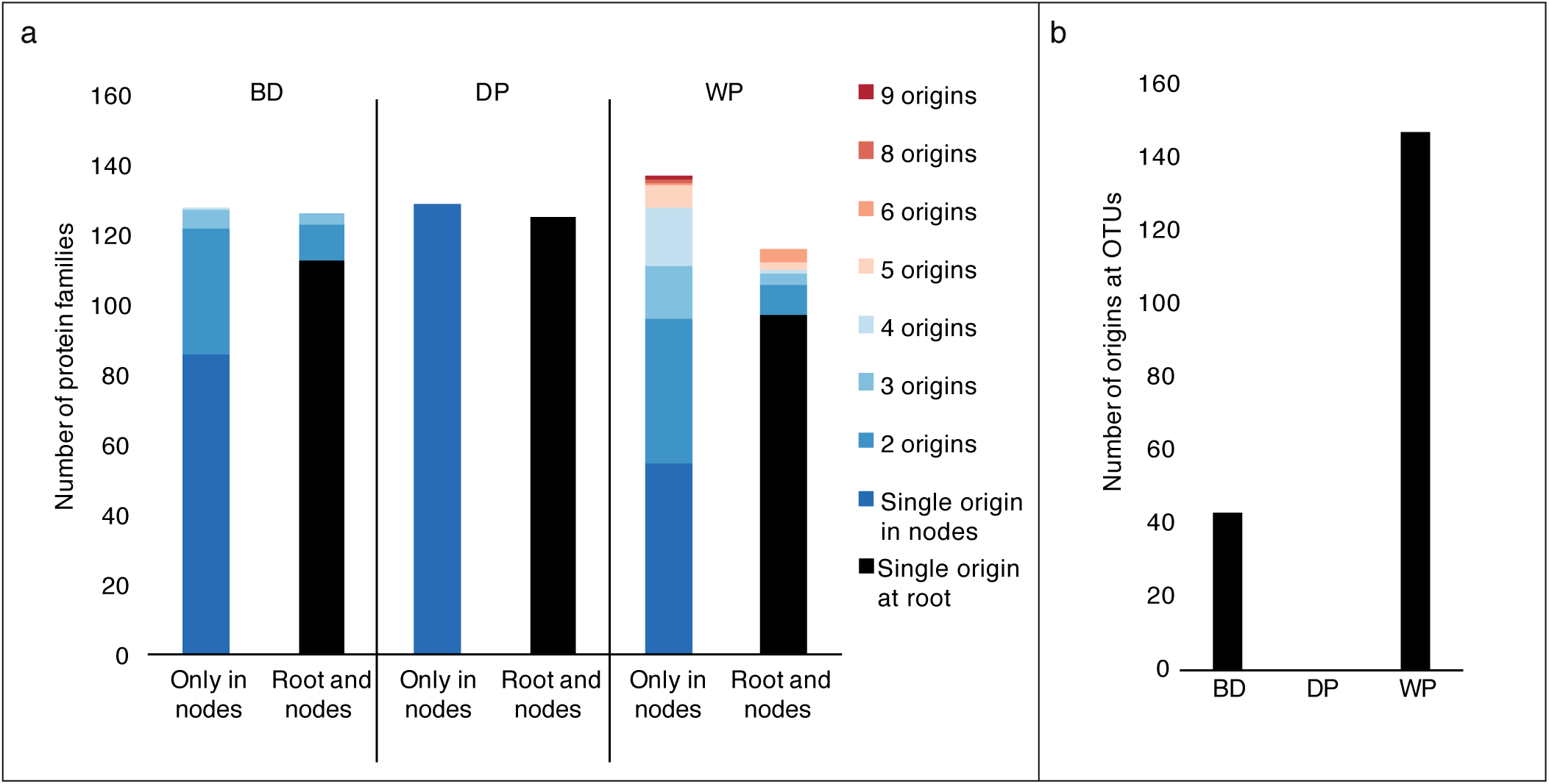
Multiple origins for the same protein families in the PL dataset calculated by Count. **(a)** Number of different gains per protein family (split by gains only in nodes or at the root and nodes) for each phylogenetic model in Count; single origins at the root are highlighted in black; a gradient from blue to red shows multiple origins for the same protein family. **(b)** Number of origins in the outer nodes of the tree for each phylogenetic model in Count.

In the 112 years since Mereschkowsky (1905) suggested that plastids arose from cyanobacteria, no one has seriously proposed a stepwise acquisition of plastid genomes. Rather, plastid endosymbiosis operates via mass acquisition of genes at the cyanobacterial origin of the organelle, followed by gene loss and transfer to the nucleus (Martin and Müller 1998; Martin and Herrmann 1998; Timmis et al. 2004; Archibald 2015). Count however delivers a result that clearly suggests "continuous" LGT into and among the members of the eukaryotic lineage in order to construct plastids “on the fly" in independent eukaryotic lineages. That is important because the central argument of Groussin et al (2016) was that Count "*supports the continuous acquisition of genes over long periods in the evolution of Archaea*". The suspicion is that Count is biased towards the inference of continuous acquisition and does not recover expected events of periodic massive gains followed by gradual differential loss even when that is the true process. Hence, this raises serious concerns about the critique by Groussin et al (2016) as their conclusions are likely a misleading outcome of the program they used, not an attribute of the data they analyzed or the evolutionary process that generated it.

Figure 2 shows the gain events calculated by the three models plotted against the reference tree. Eleven is the maximum number of gains at an OTU for the BD model (also high for WP with 17 gains) at *Pyramimonas parkeae,* a model organism for early-evolved Viridiplantae (Satjarak and Graham 2017). Wagner Parsimony places the highest number of gain events (nineteen) at *Nephroselmis olivacea,* which is considered a descendant of the earliest-diverging green algae (Turmel et al. 1999). It should be noted that all models place a considerable number of gain events at the common ancestor of Rhodophyta, Hacrobia and SAR.

Wagner Parsimony predicts the largest number of different gain events for the same protein families (Figure 3a) – eight different origins for ycf20, a family of unknown function and nine for cysT, a sulfate transporter. The BD model predicts a maximum of 4 different origins for ycf47, a poorly characterized probable protein exporter in thylakoid membranes. Strikingly, Dollo Parsimony does not predict more than one origin for any family, with only one gain event for all other 129 proteins occurring somewhere else throughout the tree. In other analyses (Martin et al. 2002) the corresponding patterns were identified as being the result of multiple independent gene losses. Both the BD and WP models predict a large number of gain events at the leaves of the reference tree – 43 and 147, respectively. (Figure 3b).

All three models in Count calculate at least one loss event per protein family for more than half of the families in the dataset (Supplemental Figure 3). However, the number of gains (LGTs or convergent gene sequence homology origin) and losses per protein family is on the same order of magnitude. This is evident on the result of the functional annotation of gain and loss events done for the PL dataset (Figure 4). We annotated 224 of the 254 families. With the exception of Dollo Parsimony for photosystem II proteins and Calvin cycle, the tree models in Count predict at least one gene gain event in all the functional categories.

**Fig. 4:**
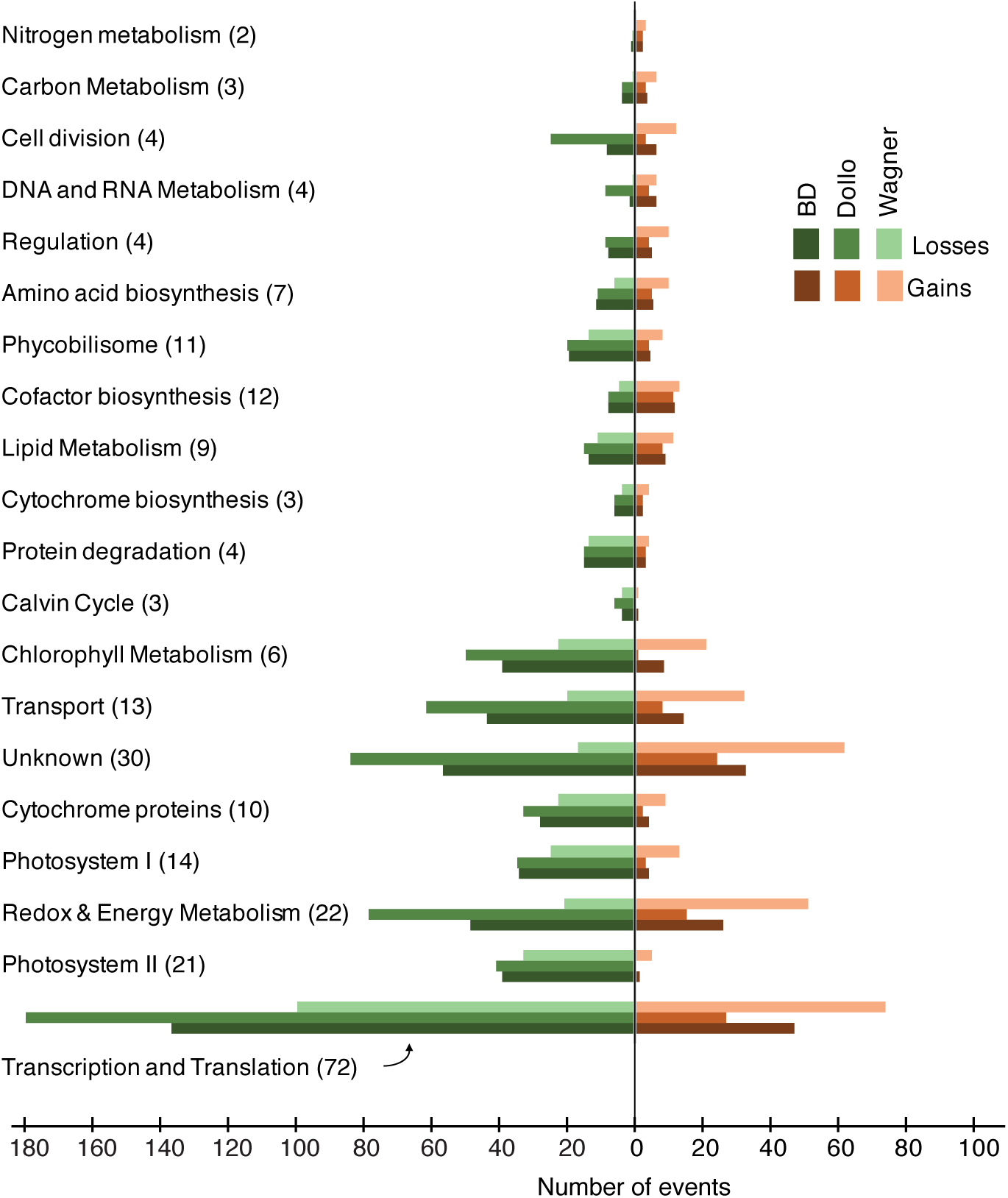
Gain and loss events for functional categories of protein families in the PL dataset. The manual annotation resulted in 20 categories listed on the y axis, sorted by the prevalence in the PAP (in parenthesis the total number of families in each category). Lost and gain events are shown on the left (greens) and right (oranges) side of the barplot, in the same scale, for the 3 different models in Count.

### The birth-and-death model of endosymbiosis events

Current views of eukaryote origin have it that eukaryotes arose from a symbiotic association between an archaeal host lineage and a mitochondrial endosymbiont (McInerney et al. 2014; Zaremba-Niedzwiedzka et al. 2017; Martin et al. 2017b) involving gene transfers from endosymbiont to host (Timmis et al. 2004; Thiergart et al. 2012). The origin of plastids entailed an additional influx of genes at the origin of the plant lineage (Ku et al. 2015). Thus, mitochondria and plastids each are currently understood to have had different, single origins, where large portion of the endosymbiont genomes entered the eukaryotic lineage. We checked the ability of the birth-and-death model from Count to recover the massive episodic gene acquisition events at the origin of eukaryotes and chloroplasts, using PAPs prepared from the EK dataset (Ku et al. 2015). The distribution of those families is shown in Figure 5, which is reproduced with permission from Ku et al. (2015).

**Fig. 5.**
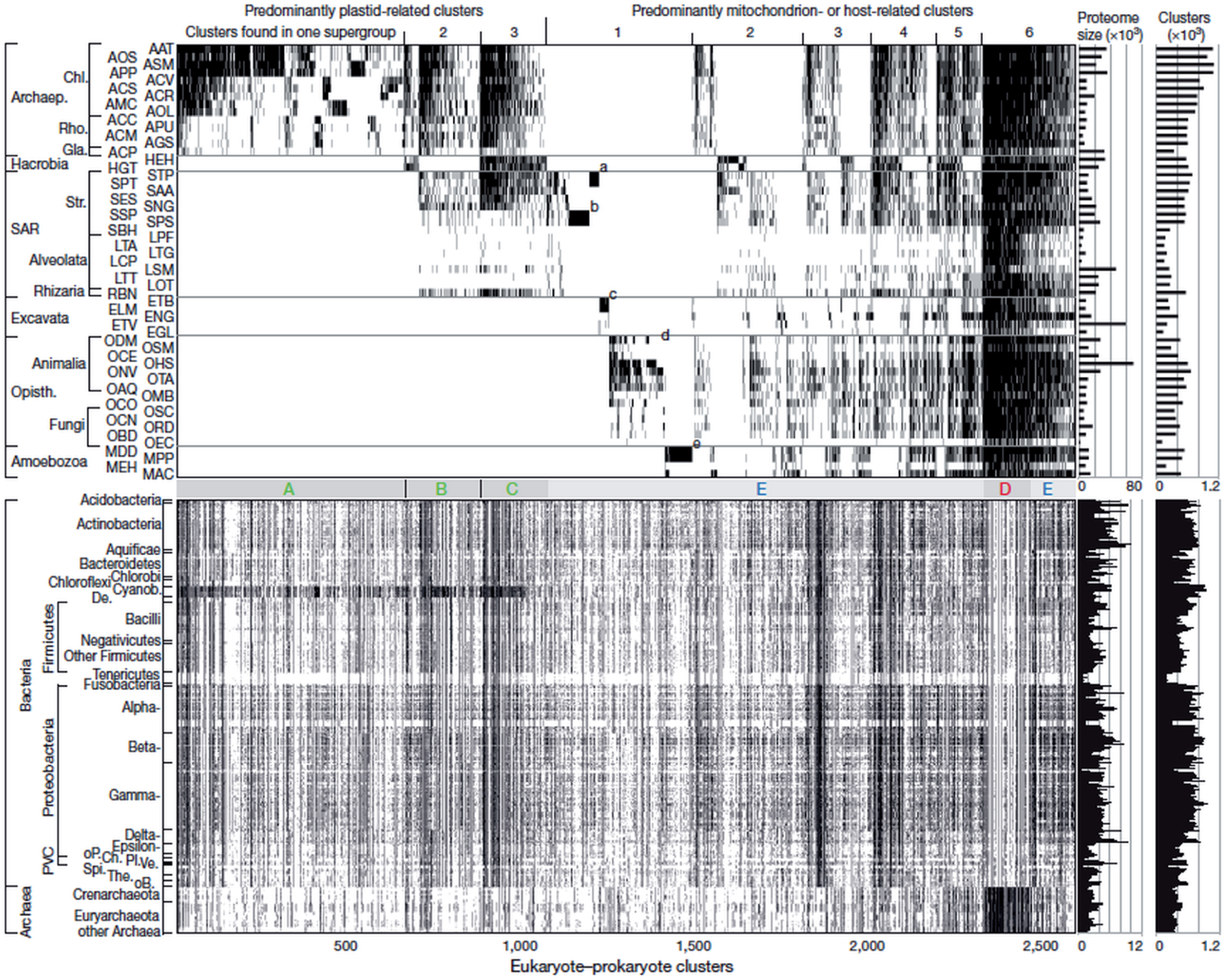
Gene distributions for eukaryotic genes. Reproduced with permission from Ku et al. (2015).

Indeed, Count’s BD model placed 1410 of all 2972 origin events for Group E proteins on the terminal edges of the phylogenetic reference tree (Figure 6a). The largest number of different gain events in a single OTU - 98 - was calculated for *Amphimedon queenslandica*, a sponge species known as a model for studying the origin and early evolution of animals (Srivastava et al. 2010). At the inner nodes of the tree the gain events were distributed almost uniformly, with only eight of the 53 inner nodes receiving no gain events with Count.

**Fig. 6:**
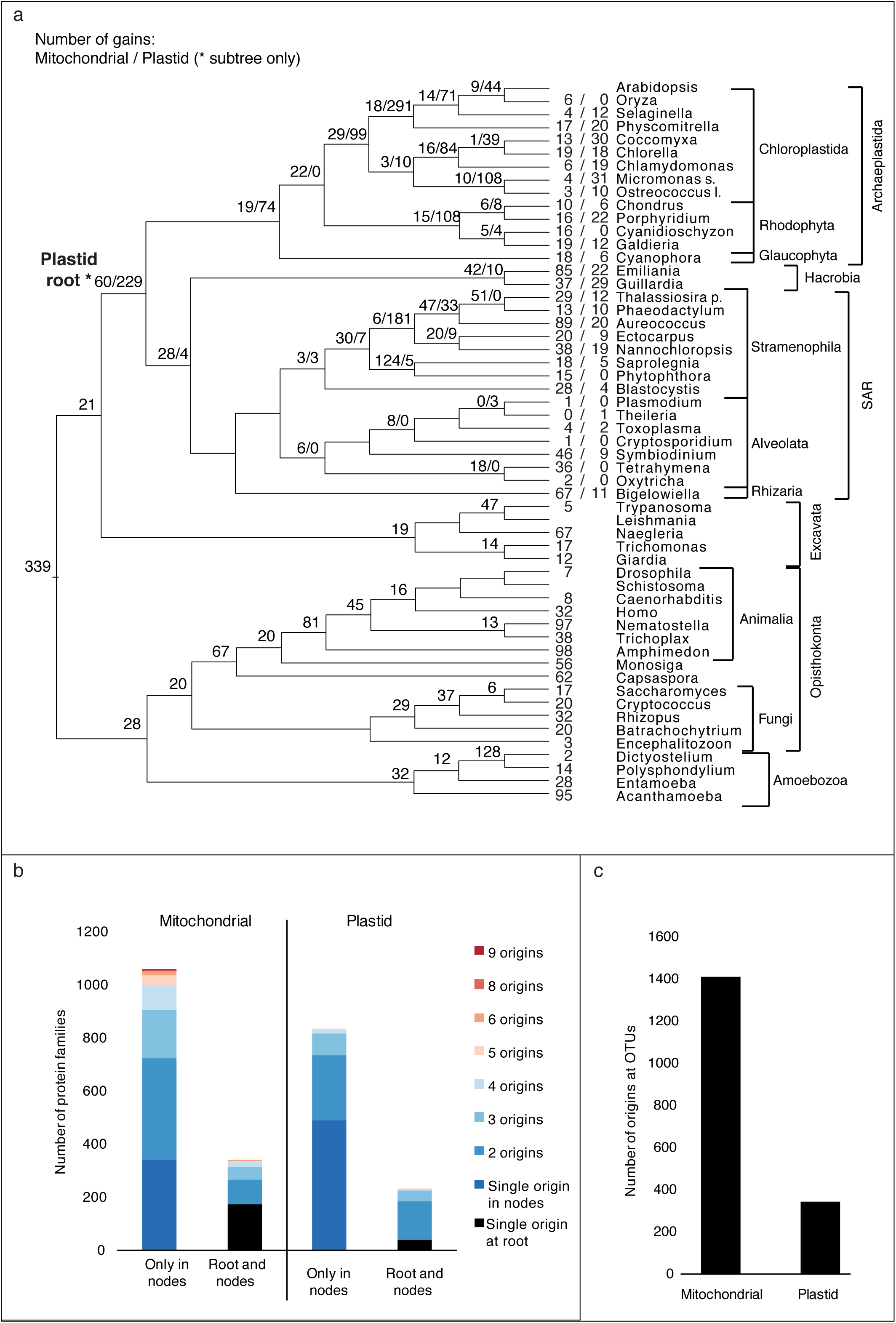
Gain events calculated by Count’s birth-and-death model for mitochondrial and plastid protein families in the EK dataset. **(a)** Reference tree for eukaryotes with mitochondrial and plastid origin events depicted at the respective nodes (separated by slashes), in this order. On the right, the 6 supergroups and the individual species (complete names in Supplemental Table 2) are shown. The root for the plastid subtree is highlighted with a star (*). (b) Number of different gains per protein family (split by gains only in nodes or at the roots of each organelle’s tree and nodes) for each phylogenetic model in Count; single origins at the root are highlighted in black; a gradient from blue to red shows multiple origins for the same protein family. (c) Number of origins in the outer nodes of the tree for each phylogenetic model in Count.

Out of 1,397 eukaryotic protein families belonging to Group E (see Figure 5) Count calculated that only 172 had a single origin at the root and no other gains anywhere else on the tree (Figure 6b). An additional 168 mitochondrial families were present at the root, however with additional origins spread throughout the tree (between one and 5 different origins). For 885 of the Group E protein families Count calculated between two and eight independent gain events (from prokaryotes via LGT or via eukaryote-eukaryote LGT). Count places a massive number of gain events at the leaves - 1410 - for the Group E protein families (Figure 6c). It is important to recall that for the 2585 genes families present in eukaryotes and prokaryotes in the data set of Ku et al. (2015), 87% show evidence for a single origin at the root using maximum likelihood methods (Ku et al. 2015). By contrast, Count reports that eukaryotes have acquired 88% of their genes independently from prokaryotes, but *from the same prokaryotic* donor each time, because otherwise the gene trees would not reflect a single origin relative to prokaryotic homologues (Ku et al. 2015). Clearly, Count does not model adequately mass acquisitions such as those incurred at endosymbiotic events that gave rise to organelles.

In the case of plastid families (Group A, B, and C in Figure 5), the genes for which are conspicuously widespread among cyanobacteria (Figure 5), Count produces the same effect: only 38 proteins out of 1060 originate once and at the root of the subtree for plastid-containing species (Figure 6a and 6b). Count attributes another 191 families to the root and with additional origins elsewhere on the tree (between two and five different origins). According to Count, eukaryotes and plastids would have been acquiring the genes for the proteins that they need to survive “on the fly”, that is via independent gains (of the same genes in independent lineages) during eukaryotic origin.

Furthermore, phylogenetic testing has shown that the vast majority of eukaryotic proteins in Group A, B, C, and E having homologues in prokaryotes are monophyletic, such that a single origin, not multiple origins, is the preferred model (Ku et al. 2015). Count does not recover that aspect of the data. Moreover, Ku et al. (2015) tested to see whether eukaryote to eukaryote LGT could account for the patchy distribution of the eukaryotic genes in Figure 5. The result was that gene evolution in eukaryotes is resoundingly vertical (Ku et al. 2015), not lateral as in prokaryotes, hence the many independent origins (LGT) that Count infers do not reconcile with the phylogenies of the proteins underlying the PAPs with which Count operates. Out of the 1,761 calculated origins of the different plastid protein families, 339, almost a fifth, were found at leaves (Figure 6c). In only nine out of the 31 inner nodes of the plastid subtree there were no gain events of plastid families. Again, for the 2585 genes families present in eukaryotes and prokaryotes in the data set of Ku et al. (2015), 87% show evidence for a single origin at the eukaryotic root using maximum likelihood methods (Ku et al. 2015).

By contrast, Count reports that plastid bearing eukaryotes have acquired 96% of their genes independently from prokaryotes, but *from the same prokaryotic* donor each time, because otherwise the gene trees would not reflect a single origin relative to prokaryotic homologues (Ku et al. 2015). Clearly, Count is doing something very unusual with PAP data in the case of mass acquisitions such as those incurred at endosymbiotic events that give rise to organelles. The same is almost certainly true for the mass acquisitions in archaea, where Count imposes a uniform process of acquisition upon the data, regardless of what the true process was.

## Discussion

LGT is important in archaea (Wagner et al. 2017). Two recent studies have indicated that in archaea, gene acquisitions from bacteria can be episodic (Nelson-Sathi et al. 2012; Nelson-Sathi et al. 2015), similar results were found for transfers at the origin of eukaryotes and at the origin of plastids (Ku et al. 2015). Groussin et al. (2016) used the results of Count (Csűrös 2010) as evidence that LGT in archaea is uniform, not episodic. We checked to see if Count could recognize loss-only as the true model. We investigated proteins encoded in plastid genomes, which were sequestered from the cyanobacterial lineage ca. 1.6 billion years ago and have been vertically inherited in eukaryotes since, except during secondary endosymbiotic events. We analyzed the three different methods for ancestral reconstruction available in Count: the birth-and-death (BD) model, Dollo Parsimony (DP), and Wagner Parsimony (WP). The results obtained show that with BD and WP, Count distributes the origin of eukaryotic protein families uniformly throughout the tree and that more than one eukaryote LGT event is often calculated for the same protein family. With DP, there are also gain events throughout the tree, although not at the leaves (OTUs) and not twice for the same family.

The results of Count would suggest a process of continuous LGT for plastids and for eukaryotes, which runs counter to data (Ku et al. 2015; Ku and Martin 2016), the standard Darwinian paradigm of eukaryote evolution (Martin 2017), and eukaryote diploid genetics (Charlesworth et al. 2017). Count has it that different eukaryotic lineages independently assembled the collections of genes that make them eukaryotic (Figure 6) and that plastids independently assembled their genomes to look like reduced cyanobacterial genomes (Figure 2 and 3). Such inferences cannot be true.

The results from Count, while unusual, can be easily explained as a consequence of the assumption of independence of gene families. Clearly this assumption is violated in the cases of acquisition and loss studied here. However, the assumption could also distort inferences made in a more general setting (Lassalle et al. 2017). There are two main, but related, effects. Firstly, the relative cost, to parsimony scores of likelihood, of LGTs are skewed. It becomes cheaper to posit separate LGTs for each gene family. Secondly, the independence of family means that the history for each gene family is inferred separately with no sharing of information across families. As each gene history is inferred using only the PAP for that family, the position of LGTs fit individually irrespective of whether they make sense in the larger context. The result is a classic case of overfitting, akin to an interpolating curve which bends and stretches to fit through every single data point.

This is not a theoretical criticism: we have shown here that this problem has real and significant impact on inference. In particular, the systematic error explains the failure of Groussin et al. (2016) to recover the patterns of archaeal LGT discovered in Nelson-Sathi et al. (2015).

The incorporation of dependence between gene families into methods like Count would be challenging both computationally and mathematically. Significant progress towards a heuristic solution has been made recently by Lassale et al. (2017). However, it could be still impossible to distinguish convincingly between different scenarios based only on PAP data, there is simply insufficient information per gene family, and it might be statistically impossible to discriminate between radically different histories. The tests implemented by Nelson-Sathi et al. (2015) lacked the statistical power of full likelihood-based methods (Yang et al. 2007), but on the other hand made few assumptions on the process of LGT accumulation, gaining some robustness in turn.

## Acknowledgments

This work was supported by grants from the ERC (666053) and the Volkswagen Foundation (93 046) to WFM, the German Israeli Foundation [(I-1321-203.13/2015] to WFM and EHC, the Open University of Israel Research fund (504735) to EHC, the Department of Science and Technology INSPIRE Faculty Award (DST/INSPIRE/04/2015/002935) to SN-S and the New Zealand Bio-Protection Research Centre 2017 PI funding allocation to PJL.

